# KSHV genome harbors both constitutive and lytically induced enhancers

**DOI:** 10.1101/2024.01.30.577957

**Authors:** Nilabja Roy Chowdhury, Vyacheslav Gurevich, Meir Shamay

**Author notes:** Corresponding author Meir Shamay **Email:**. **Author Contributions:** Conceived and designed the experiments: N.R.C., V.G., M.S. Performed the experiments: N.R.C., Analyzed the data: N.R.C., V.G., M.S. Generated tools & reagents: N.R.C., V.G., Wrote the paper: N.R.C. and M.S. **Competing Interest Statement**: The authors declare no competing financial interests.

## Abstract

Kaposi’s sarcoma-associated herpesvirus (KSHV) belongs to the gamma-herpesvirus family and is a well-known human oncogenic virus. In infected cells, the viral genome of 165 kbp is circular DNA wrapped in chromatin. The tight control of gene expression is critical for latency, the transition into the lytic phase, and the development of viral-associated malignancies. Distal cis-regulatory elements (CRE), such as enhancers and silencers, can regulate gene expression in a position and orientation-independent manner. Open chromatin is another characteristic feature of enhancers. To systematically search for enhancers, we cloned all the open chromatin regions in the KSHV genome downstream to the luciferase gene and tested their enhancer activity in infected and uninfected cells. A silencer was detected upstream of the latency promoter (LANAp). Two constitutive enhancers were identified in the K12p-OriLyt-R and ORF29 Intron region, where ORF29 Intron is a tissue-specific enhancer. The following promoters: OriLyt-L, PANp, ALTp, and the Terminal Repeats (TRs) acted as lytically induced enhancers. Expression of the Replication and Transcription Activator (RTA), the master regulator of the lytic cycle, was sufficient to induce the activity of lytic enhancers in uninfected cells. We propose that the TRs that span about 24 kbp region serve as a “viral super-enhancer” that integrates the repressive effect of the latency protein LANA with the activating effect of RTA. Utilizing CRISPR activation and interference techniques, we determined the connections between these enhancers and their regulated genes. The silencer and enhancers described here provide an additional layer to the complex gene regulation of herpesviruses.

**Importance:** In this study, we performed a systematic functional assay to identify cis-regulatory elements within the genome of the oncogenic herpesvirus, Kaposi sarcoma-associated herpesvirus (KSHV). Similar to other herpesviruses, KSHV presents both latent and lytic phases. Therefore, our assays were performed in uninfected cells, during latent infection, and under lytic conditions. We identified two constitutive enhancers, where one seems to be a tissue-specific enhancer. In addition, four lytically induced enhancers, which are all responsive to the Replication and Transcription Activator (RTA), were identified. Furthermore, a silencer was identified between the major latency promoter and lytic genes locus. Utilizing CRISPR activation and interference techniques, we determined the connections between these enhancers with their regulated genes. The terminal repeats spanning a region of about 24 kbp, seem like a “viral super-enhancer” that integrates the repressive effect of the latency protein LANA with the activating effect of RTA to regulate latency to lytic transition.

## Introduction

Kaposi’s Sarcoma-associated Herpesvirus (KSHV/HHV8) belongs to the γ-herpesvirus family and is one of the well-known human oncogenic viruses. KSHV is the etiologic factor for Kaposi’s Sarcoma (KS), and is associated with Primary Effusion Lymphoma (PEL), Multicentric Castleman’s Disease (MCD) and KSHV-associated Inflammatory Cytokine Syndrome (KICS) (1). The large double-stranded DNA genome of ∼165kb encodes numerous Open Reading Frames (ORFs) and multiple non-coding RNAs (ncRNA). Like other herpes viruses, KSHV presents both latent and lytic modes of infection. Only a handful of viral genes, including LANA, vFLIP, vCyclin, Kaposin, and 12 micro-RNA (miRNA) are expressed during latency (2). In sharp contrast, during the lytic/productive phase, almost ninety viral proteins are expressed, including proteins responsible for DNA synthesis, virion assembly, and release (2). The tight control of gene expression is critical for the maintenance of latency, the transition into the lytic phase, the correct order of lytic gene expression, as well as the development of viral-associated malignancies.

Upon infection, the KSHV viral genomes circularize and reside in the nucleus of the infected cells as large plasmids known as episomes (3–5). Initially, these episomes are wrapped by histones with open chromatin marks, such as tri-methylated Histone 3 Lys4 (H3K4Me3) and acetylated Lys27 (H3K27Ac) (6, 7). Later, following the recruitment of the Polycomb Repressive Complex (PRC), an increase in H3K27Me3 and H2AK119ub histone repressive marks are observed (7, 8). The Latency-associated Nuclear Antigen (LANA) protein plays an important role in maintaining the episomes by binding to multiple LANA Binding Sites (LBS) in the viral Terminal Repeats (TRs) (9). Each TR is a high GC repeats region of 801 bp containing three LBSs (LBS1, LBS2 and LBS3) (10–13). LANA also binds additional sites on the KSHV genome, including the promoter of the Replication and Transcription Activator (RTA/ORF50) (14, 15). Induction of lytic reactivation can be achieved by combined treatment of KSHV-infected cells with Phorbol esters and Sodium Butyrate (NaB), which can turn on the expression of RTA and promote histone acetylation (16, 17). RTA expression is both sufficient and required for the initiation of lytic reactivation, as it binds and activates multiple lytic gene promoters (18–22). Many lytic promoters are RTA-responsive regions, including the two viral lytic replication origins (OriLyt-L and OriLyt-R), and Polyadenylated nuclear (PAN) RNA promoter (23–26). Therefore, by binding and repressing the RTA promoter, LANA plays another crucial role in maintaining the viral latency (27–29).

Successful expression of a gene is precisely controlled by the gene promoter and its distal cis-regulatory elements (CREs), such as enhancers and/or silencers (30). This control involves the recruitment of cohesion complexes, CCCTC-binding factor (CTCF), Yin-Yang 1 (YY1), the mediator complex, specific and general transcription factors, RNA polymerase II (PolII), and chromatin remodelers and modifiers (31). CREs can regulate distal promoters by interacting with the respective promoter through DNA loop formation (30). In contrast to promoters, enhancers can affect gene expression in a position- and orientation-independent manner (32).

Although enhancers and promoters were regarded as different entities, both past and recent studies have shown that bi-directional active promoters can also serve as enhancers (33, 34), known as ‘E-promoters’ (35). The E-promoters, like the conventional enhancers, exhibit H3K27Ac but also show H3K4Me3 instead of H3K4Me1 (36). Silencers are CREs that function opposite to enhancers, i.e., they repress the expression of genes by binding with repressors and interacting physically with respective gene promoters (37, 38). Silencers are located at a variable distance of 200-2000bp from the respective gene promoters (39, 40). Moreover, there are reports of dual-function CREs, where the same sequence can function as both an enhancer and a silencer in different cellular contexts (41, 42).

When two regulatory elements are in close proximity, distinguishing which is the enhancer and which is the promoter, in particular within a condensed viral genome, can be challenging. While H3K4Me1 histone marks might assist (43), recent studies suggest that enhancers can also be marked with variable degrees of H3K4Me3 marks (44, 45). Therefore, the best way to determine enhancers and silencers is by a functional assay that exploits their ability to affect gene expression while being situated downstream to the promoter. Self-Transcribing Active Regulatory Regions sequencing (STARR-seq) is a powerful assay to determine enhancers genome-wide, where downstream enhancers induce a weak promoter to facilitate mRNA transcription (46). Formaldehyde Assisted Isolation of Regulatory Elements sequencing (FAIRE-seq) serves as a robust method to detect open chromatin regions of a given genome harboring CREs (47, 48). FAIRE-seq in KSHV latently infected cells identified 24 open chromatin regions (49). In the present study, we aimed to systematically search for enhancers and silencers in the KSHV genome. Therefore, all FAIRE-seq regions were cloned in a STARR-seq library plasmid downstream to a weak promoter controlling the luciferase expression. Transfection of this KSHV-FAIRE-STARR library into KSHV-infected and un- infected cells revealed 6 viral enhancers. ORF29 Intron and K12p-OriLyt-R were identified as constitutive enhancers, while OriLyt-L, PAN promoter, ALT promoter and the Terminal Repeats (TR) were identified as RTA inducible lytic enhancers. We also found that a region within the LANA promoter (LANAp) acts as a silencer element in the KSHV genome. Interestingly, the TR, which acts as a lytically inducible enhancer, harbors a putative binding motif for RTA at the LBS1. Mutation of this site in the TR significantly diminished RTA-mediated TR enhancer activity. It was also observed that RTA can compete with LANA to induce TR enhancer activity. To reveal their functional significance, viral enhancers were targeted with dCas9 fused to interfering/repressing (CRISPRi) or activating (CRISPRa) transcriptional domains. This resulted in disruption of the viral transcriptional repertoire in both latency and lytic phases. In summary, our findings reveal that the KSHV genome harbors a silencer and several constitutive and lytically induced enhancers that control viral gene expression during latency and lytic phases. This study also provides insights into the differential viral gene expression in different types of KSHV-infected cells and tissues.

## Materials and Methods

### Cell Lines

BCBL-1 cells were maintained in RPMI-1640 medium supplemented with 20% Fetal Bovine Serum (FBS), L-glutamine (2mM, Gibco), Penicillin-Streptomycin (100IU/ml and 100ug/ml respectively, Gibco), Sodium-pyruvate (1mM, Gibco) at 37°C under 5% CO2 atmosphere. iSLK, iSLK.219, SLK.219, HEK293, HEK293T and HEK293.219 cells were grown in DMEM medium supplemented with 10% Fetal Bovine Serum (FBS), with the same supplements. HCT116 cells were maintained in McCoy’s 5A medium supplemented with 10% Fetal Bovine Serum (FBS), with the same supplements. The cell lines infected with rKSHV.219 were maintained in 1ug/ml Puromycin selection. Unless stated otherwise, iSLK cells were treated with 1ug/ml Doxycycline to induce RTA expression in respectively mentioned experiments.

### Lytic induction of KSHV and generation of free virions

SLK.219 and HEK.219 cells were treated with 20ng/ml Phorbol Ester (TPA) and 1.25mM Sodium Butyrate (NaB) for reporter and CRISPR experiments. iSLK.219 cells were induced with 1ug/ml Doxycycline and 1.25mM NaB for lytic reactivation. BCBL-1 cells were treated with the same concentrations of TPA and NaB to generate KSHV virions. Briefly, BCBL-1 cells were treated with TPA and NaB on the first day, followed by only TPA for four more days. Centrifugation was done thereafter @ 600g for 5-7 minutes at 4°C. The supernatant was filtered with 0.45um filters. The filtrate containing the free virion was centrifuged @ 15000g for 10 hours at 4°C. The obtained virion pellet was resuspended in 400ul ice-cold Dulbecco’s Phosphate-Buffered Saline solution (DPBS).

### Isolation of KSHV viral DNA

To obtain pure KSHV virion DNA, the DPBS resuspended virion pellet was treated with RNase free DNase Set (Qiagen, Cat. No. 79254) which was neutralized with 10mM Ethylenediamine Tetra-acetic Acid (EDTA) pH 8 at 65°C for 45 minutes. DNA isolation was then performed using DNeasy Blood and Tissue Kit (Qiagen, Cat. No. 69506) following the manufacturer’s protocol.

### Generation of KSHV FAIRE-STARR library

Open chromatin regions in the KSHV genome were identified from the study by Hilton et al., 2013 (49) and extended by ∼200bp on each side to design primers for Gibson assembly (**Table S1**). PCR was performed using 21 pairs of primers (**Table S2**) using KSHV viral DNA as a template. Purified amplicons were cloned into STARR-seq luciferase reporter vector within the XbaI and FseI sites using Gibson assembly master mix (NEB Cat. No. E5510). The STARR-seq luciferase reporter (vector_ORI_empty) was a gift from Alexander Stark (Addgene plasmid # 99297 ; http://n2t.net/addgene:99297 ; RRID:Addgene_99297). KSHV TRs were cloned from TR7-LucP vector, a gift from Rolf Renne (10), into BamHI and SalI sites of Addgene #99297 plasmid using T4 DNA ligase (NEB, Cat. No. M0202).

### Transfection of FAIRE-STARR library into Cell Lines

Cell lines used in reporter assays were co-transfected with KSHV-FAIRE-STARR plasmids and internal control pGL4.74 (Renilla) plasmid using PolyJet (Signagen, Cat. No. SL100688) with the exception of BCBL-1 cells that were transfected with FuGENE HD transfection reagent (Promega, Cat. No. E2311,). Competition reporter assays were performed with co-transfection of the TR7 reporter plasmid with or without LANA expression plasmid (DY52). Reporter signals were read with Multilable Microplate Reader (TECAN Infinite M1000).

### Generation of CRISPRi and CRISPRa plasmids and lentiviral transduction

HEK293T cells were transfected with pLX-TRE-dCas9-KRAB-MeCP2-BSD plasmid, a gift from Andrea Califano (Addgene plasmid #140690; http://n2t.net/addgene:140690; RRID: Addgene_140690) (50) for CRISPRi or ScFv-2ERT2-VPH, a gift from Yu Wang (Addgene plasmid #120556; http://n2t.net/addgene:120556; RRID: Addgene_120556) (51), pHRdSV40- dCas9-10xGCN4_v4-P2A-BFP (Addgene plasmid # 60903; http://n2t.net/addgene:60903; RRID: Addgene_60903), a gift from Ron Vale (52) plasmids for CRISPRa with specific sgRNA (**Table S2**) targeting KSHV enhancers and PSPAX2 lentiviral expression plasmid, a gift from Didier Trono (Addgene plasmid #12260; http://n2t.net/addgene:12260; RRID: Addgene_12260), to generate lentiviral particles. SLK.219 cells were transduced with lentiviruses supplemented with 8ug/ml polybrene. The cells were selected with 1ug/ml Blasticidin (for CRISPRi transduced cells selection) or 700 ug/ml G418 and 600 ug/ml Hygromycin (for CRISPRa transduced cells selection) for 7 days. For CRISPRi, cells were induced with 1ug/ml Doxycycline for 2 days, followed by 20ng/ml TPA for lytic induction. For CRISPRa, cells were induced with 200 nM 4-hydroxytamoxifen (4-OHT) for 72 hours.

### RNA-sequencing

RNA from the CRISPRi and CRISPRa cells was purified by RNeasy Mini kit (Qiagen, #74106) according to the manufacturer’s instructions. The integrity of the isolated RNA was tested using the Agilent TS HSRNA Kit and Tapestation 4200 at the Genome Technology Center at the Azrieli Faculty of Medicine Bar-Ilan University. 1000 ng of total RNA were used for mRNA enrichment by using NEBNext mRNA polyA Isolation Module (NEB, #E7490L), and libraries for Illumina sequencing were performed using the NEBNext Ultra II RNA kit (NEB, #E7770L). Quantification of the library was performed using dsDNA HS Assay Kit and Qubit 2.0 (Molecular Probes, Life Technologies), and qualification was done using the Agilent D1000TS Kit and Tapestation 4200. 500 nM of each library was pooled together and then diluted to 4nM according to NextSeq manufacturer’s instructions. 1.5pM was loaded onto the Flow Cell with 1% PhiX library control. Libraries were sequenced by Illumina NextSeq 550 platform with single-end reads of 75 cycles according to the manufacturer’s instructions.

### Bioinformatic Analysis

FASTq files obtained from the Illumina NextSeq 550 platforms were concatenated for the three biological repeats using the Terminal program of MacOS Sonoma 14.2 to increase the alignment hits with the KSHV genome assembly (Accn No. NC_009333, RefSeq assembly GCF_000838265.1). The concatenated files were analysed using the SeqMan NGen tool (Version 17.6. DNASTAR. Madison, WI). Briefly, adaptor trimming was performed for all the .fastq files. The sequencing files were then aligned with the KSHV genome (Accn no. NC_009333, Ref Seq assembly GCF_000838265.1) using BowTie2 (53). ArrayStar tool (Version 17.6. DNASTAR. Madison, WI) and GraphPad Prism (v10) were used to quantitate and plot the heatmap of the analyzed RNAseq data. To develop the lines between the enhancers and respective ORFs of the viral genome, UCSC genome browser (54) was used.

## Results

### Functional assays revealed Inducible and Constitutive enhancer elements in the KSHV genome

Enhancers present different histone marks depending on their activation status. However, they consistently maintain open and accessible chromatin, which can be detected by FAIRE-seq (47). To systematically search for potential enhancers in the KSHV genome, we utilized the open chromatin regions previously identified by FAIRE-seq (49) in a functional assay (**Fig. 1A & B**). These sequences (mentioned in **Table S1**) were cloned downstream to the luciferase gene of a reporter plasmid with a weak promoter to construct the KSHV-FAIRE-STARR library (**Fig. 1C**). Some of the adjacent open chromatin regions were used in one clone, this is why we eventually had 21 clones. To maintain a consistent length of ∼1000-1200 bp, same as STARR-seq library, 200bp were added both upstream and downstream from the KSHV genome surrounding the respective FAIRE regions (**Fig. 1C**). The KSHV-FAIRE-STARR library was transfected into a KSHV-positive PEL cell line (BCBL1) and luciferase assay was performed during latency and lytic cycles. A luciferase reporter under RTA promoter was used as a positive control for lytic induction, and the Renilla reporter plasmid served as an internal control for transfection efficiency. The activity of the putative enhancers was determined relative to the same plasmid for the library that contained a non-functional genomic region, as previously used for STARR-seq (55).

**Figure 1:**
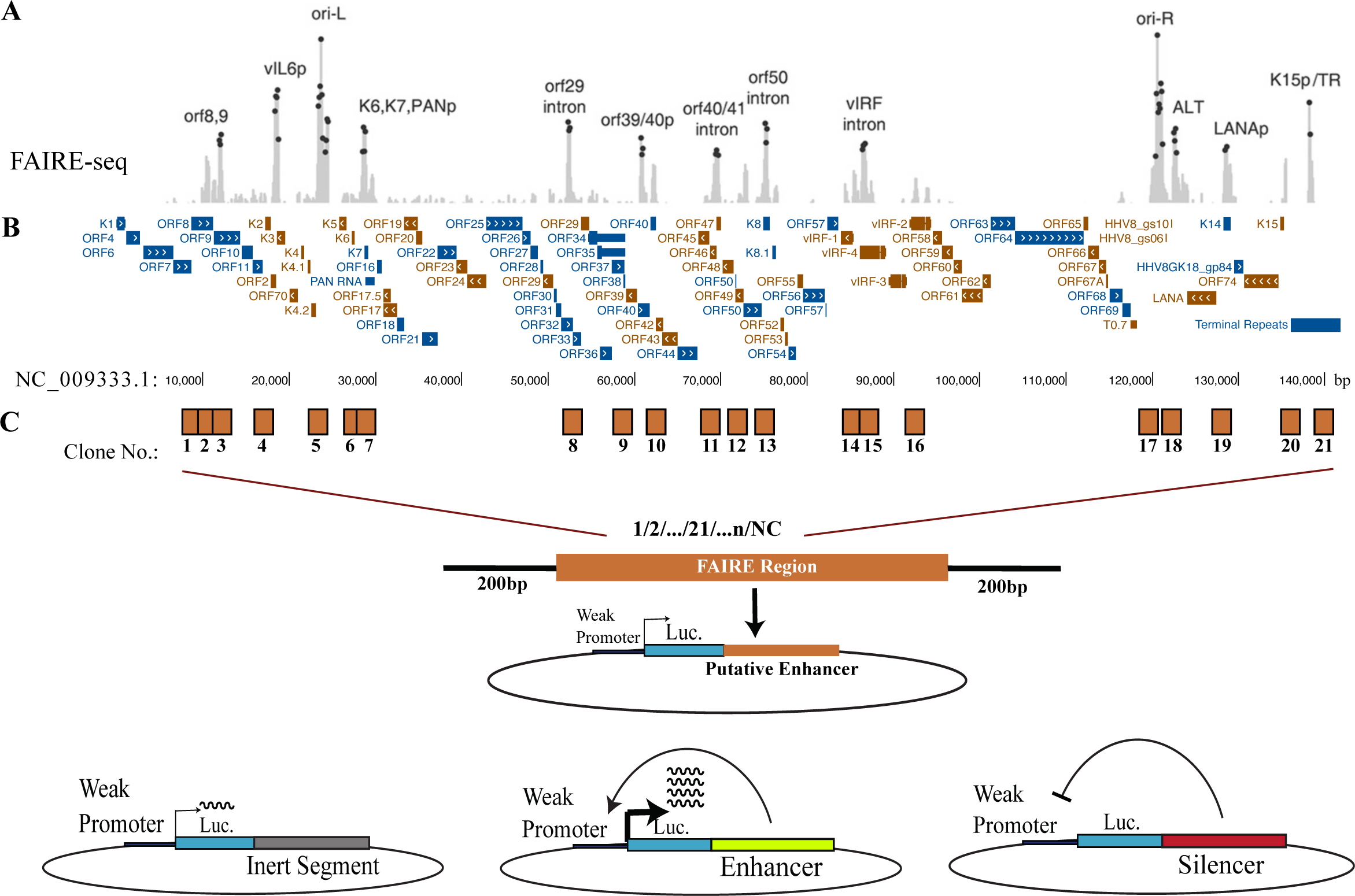
**Schematic illustration of the research strategy. (**A) FAIRE-seq peaks of the KSHV genome were adopted from Hilton et al (49). (B) Reference KSHV genome (NC_009333) schema was generated with UCSC genome browser(54). (C) Locations of the open chromatin regions identified by Hilton et al (49) and their respective clone numbers in our study. Cloning strategy and expected outcome for functional enhancers and silencers.

Reporter assays conducted on transfected BCBL1 cells revealed that several regions, including OriLyt-L (region 5), PAN promoter (region 6), K12p/OriLyt-R (region 17.1), OriLyt-R/ALTp (region 17.2) and the Terminal Repeats (region 21) exhibited significantly higher reporter signals (at least ∼2.5 fold) compared to the negative control (**Fig. 2A**). Several regions, including 5, 6, 17.2, and TR possess enhancer activity specifically in lytically induced cells, while region 17.1 exhibited activity even during latency. Enhancers are known to play an important role in tissue-specific gene regulation (56). Therefore, we were interested in testing our library in a different cell type infected with KSHV. The KSHV-FAIRE-STARR library was introduced into iSLK epithelial cells infected with rKSHV.219 (iSLK.219) and possesses a Doxycycline (Dox) inducible RTA expression system for efficient lytic induction (**Fig. 2B**). This revealed two additional enhancers: ORF29 Intron (region 8) and an intragenic region of K8.1 (region 13). In summary, in both experiments, regions 5, 6, 17.2, 21, and 13 (specific to iSLK.219) displayed enhancer activities upon lytic reactivation. This suggests that these regions harbor ‘poised’ enhancers that became ‘active’ during viral lytic reactivation due to the recruitment of certain viral immediate early (IE) proteins, along with other lytic proteins and associated transcription factors. Interestingly, region 17.1 and region 8 (in iSLK.219) showed enhancer activity even without lytic reactivation, indicating their ‘constitutive’ nature. The fact that ORF29 Intron (region 8) only acted as an enhancer in iSLK.219 (epithelial) cells but not in BCBL-1 (lymphoma B) cells, suggests that this enhancer has no activity in B-cells and fits the notion of a tissue-specific enhancer.

**Figure 2:**
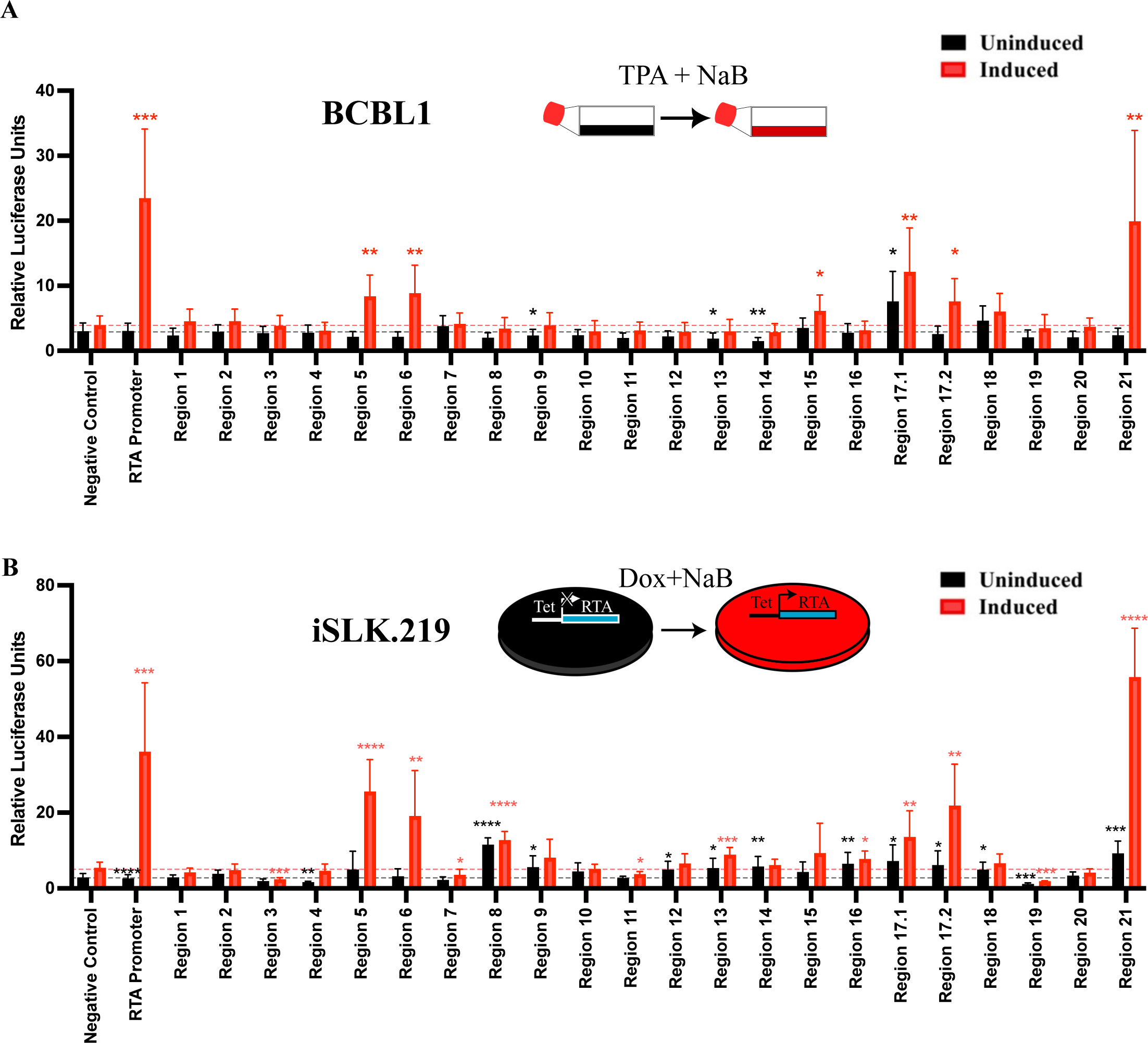
**Functional assays reveal both constitutive and lytically induced enhancers within the KSHV genome**. (A) The KSHV-FAIRE-STARR library was transfected in BCBL1 cells and luciferase assay was performed. Cells were left untreated following the transfection (uninduced), or were treated with TPA and NaB for 24 h. (B) iSLK.219 cells were transfected same as in A and induced with 1μg/ml Doxycycline and 1.25mM Sodium Butyrate for 24 hours, followed by only Doxycycline (1μg/ml) induction for another 24 hours before collecting the lysates for dual luciferase reporter assays. Relative Luciferase units represent the luciferase units that were normalized with a renilla control. Black bars indicate uninduced cells and red bars are the induced counterpart. Data represent Mean ± S.D. of n=3 biological replicates in 3 experiments. *P<0.05; **P<0.01; ***P<0.001; ****P<0.0001.

Enhancers have the ability to regulate the expression of their target genes in a position and orientation-independent manner. To investigate this further, we cloned the two most potent enhancers, OriLyt-L (region 5) and ORF29 Intron (region 8), in two different orientations and assessed their enhancer activity. As expected, they behaved as enhancers in both orientations upon lytic induction, although the inverse orientation of clone 5 presented lower activity (**Fig. S1**). Collectively, these findings indicate that the KSHV genome contains a variety of inducible, constitutive, and tissue-specific enhancer elements.

### KSHV lytic enhancers are regulated by RTA

RTA is the master regulator protein of KSHV lytic transcription. Interestingly, ChIP-seq analysis for RTA revealed its association with all the ‘open chromatin’ regions identified by FAIRE-seq regions in the KSHV genome (57, 58). Therefore, we asked whether these enhancers are responsive to RTA. To explore this, iSLK cells (uninfected) were transfected with the KSHV-FAIRE-STARR library, and the cells were treated with Dox. Expression of RTA alone was sufficient to activate all the lytically induced enhancers (**Fig. 3A**). Moreover, the reporter signals were much higher in this case, suggesting that in infected cells, a viral factor might repress these elements to contradict RTA activity. We noticed that the vIRF1 intragenic region (region 14) acted as a constitutive enhancer in uninfected iSLK.

**Figure 3:**
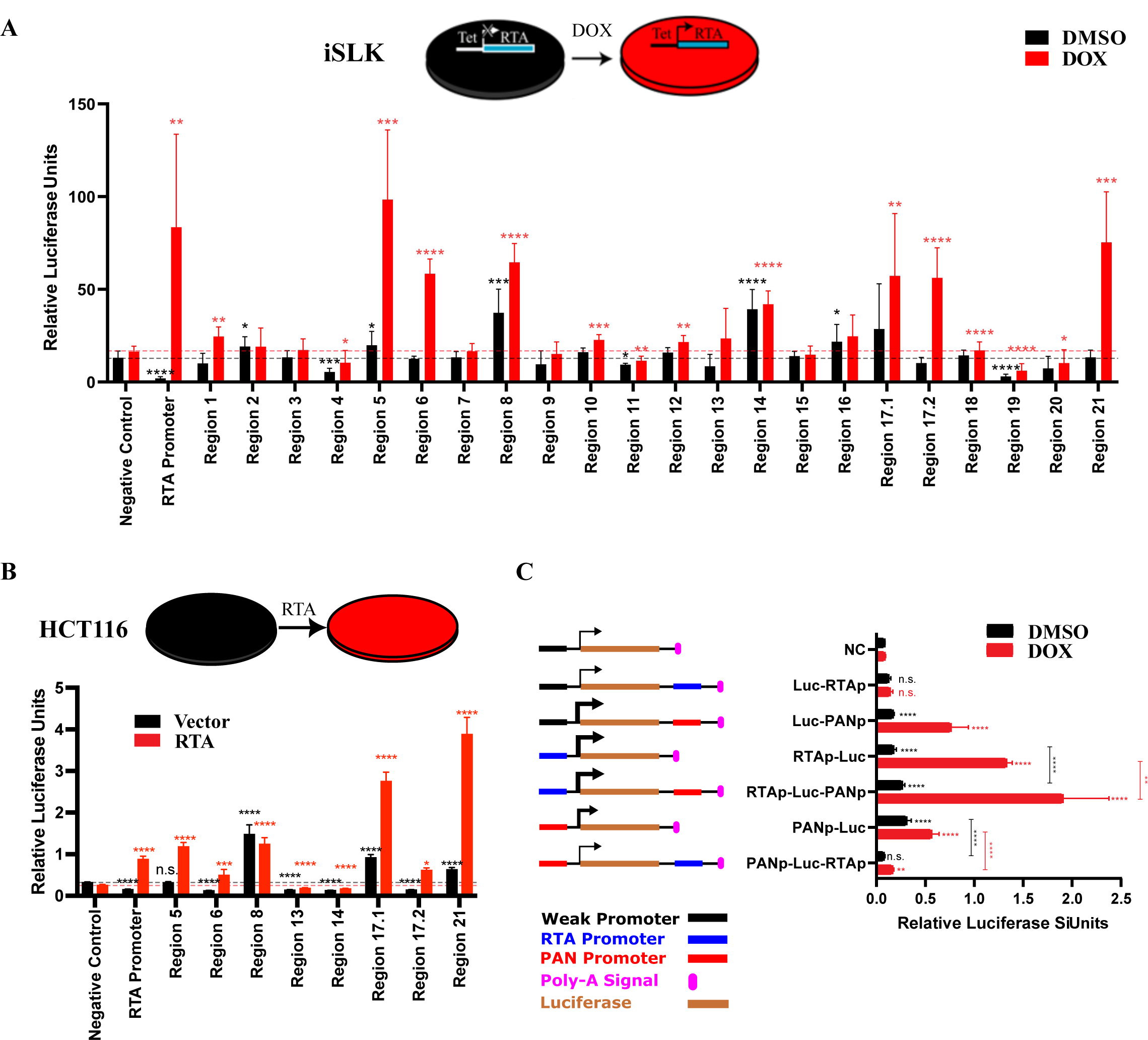
**KSHV genomic enhancers are k-RTA responsive**. (A) iSLK cells were transfected with the KSHV-FAIRE-STARR library and treated with 1μg/ml Doxycycline for 24 h before collecting the lysates for dual luciferase reporter assays. Black bars indicate uninduced cells and red bars are the induced counterpart. (B) HCT116 cells were transfected with putative enhancers with or without expression vector for RTA, and luciferase assays were performed. (C) Schematic illustrations of RTAp and PANp in promoter and enhancers position. Reporter plasmids were transfected into iSLK cells as indicated on the left and treated with 1μg/ml Doxycycline for 24 h before collecting the lysates for dual luciferase reporter assays. Data represent Mean ± S.D. of n=3 biological replicates in 3 experiments. *P<0.05; **P<0.01; ***P<0.001; ****P<0.0001.

One concern in STARR-seq is that transfection of plasmid DNA into various cell lines may provoke host interferon 1 (IFN-I) response (55) which might lead to false-positive enhancer identification. To address this issue, the STARR library is also transfected into a cell line with a non-functional interferon response, such as HCT116 (59). Transfection of all the viral enhancers from the aforementioned experiments into HCT116 cells with or without RTA expression vector revealed that all the enhancers; OriLyt-L (region 5), PANp (region 6), ORF29 Intron (region 8), K12p/OriLyt-R (region 17.1), OriLyt-R/ALTp (region 17.2) and TRs (region 21) were still active in HCT116 (**Fig. 3B**). The only exception was vIRF-1 intragenic region (region 14) construct that did not show any reporter activity in HCT116 as opposed to iSLK, suggesting this is a IFN-1 responsive enhancer. Altogether, these data indicate that all the KSHV lytically induced enhancers are RTA-responsive.

Our functional assay identified PANp as a lytic enhancer. A previous study detected strong interaction/looping between PAN promoter and RTA promoter (58). To further explore the functional relations between these two promoters, we cloned PANp downstream to RTAp, and vice-versa (**Fig. 3C**). These reporter plasmids were transfected into iSLK, and cells were treated with Dox to express RTA. This set of experiments clearly shows that PANp can function both as a promoter and an enhancer that can enhance transcription from RTAp. In contrast, RTAp can only serve as a promoter. These experiments indicate that RTAp is only a promoter, while PANp serves both as a promoter for PAN RNA and a lytic enhancer.

### RTA competes with LANA to induce TR-enhancer activity

LANA binds to the terminal repeats (TRs) to ensure the replication and tethering of the KSHV episome to the host chromosome during latency (5, 10, 60–63). It is well known that LANA can repress transcription in luciferase assays with TR (64). A commonly used reporter is 7xTR_Luc, where seven TRs are cloned downstream of the luciferase reporter gene. LANA’s ability to repress transcription in this context already suggested that TR may serve as an enhancer element (64). Our observations further suggest that RTA, in addition to LANA, can regulate the TR enhancer element, but in the opposite direction, activation instead of repression. The result that the TR reporter in our KSHV-FAIRE-STARR library (clone no. 21) can be activated by Dox alone in uninfected cells (iSLK) but not in infected cells (iSLK.219) supports the notion that infected cells express a protein that can compete with RTA for TR activity (**Fig. 4A&B**). However, even in infected cells, RTA can activate the TR if additional lytic inducers, such as sodium butyrate (NaB), are added (**Fig. 4A**). This suggests that in infected cells, higher expression levels of RTA are needed. Reporter signal for TRs increased ∼5.6-fold in iSLK.219 and ∼6-fold in iSLK cells post-induction (**Fig. 4A** and **Fig. 3**, region 21) confirming that TRs are RTA-inducible enhancers. Interestingly, ChIP-seq analysis for RTA revealed its association with the TR (65). However, to date, no specific RTA binding site on TRs has been reported. To search for common motifs between the identified enhancers, we conducted a motif search using the MEME suite(66). Although this led to the detection of 5 common motifs, none of them were found on TRs (**Fig. S2A**). This could be due to significant dissimilarities between TRs and other RTA inducible enhancers. We then examined the consensus motifs generated from the MEME suite (66) and compared them with the RTA binding motif using known RTA binding sites (57) in the MEME suite(66). A multiple motif sequence alignment using WebLogo3 (67) predicted an RTA binding motif on the TRs (**Fig. 4C**). Remarkably, this motif had striking similarities with LANA binding motifs on the human chromosome (68–70). This led us to question if the RTA binding motif is present on the LANA binding sites. Surprisingly, the particular MEME-predicted RTA-LANA sequence motif was found in TR LBS1 (**Fig. 4D**). To test if this motif is essential for the ability of RTA to activate the TR, a region of 80 bp covering LBS-1 to 3 of both wt and a mutant with three nucleotide substitutions were used in luciferase reporter assay (**Fig. 4E**). Treatment of iSLK with Dox to express RTA resulted in activation of the wt TR, but not the mutant one. This indicates that the RTA binding site within LBS1 is essential for the induction of the TR enhancer activity by RTA. The fact that wt TR contains only one TR repeat as opposed to the seven TR repeats in clone 21, may explain its relatively low enhancer activity.

**Figure 4:**
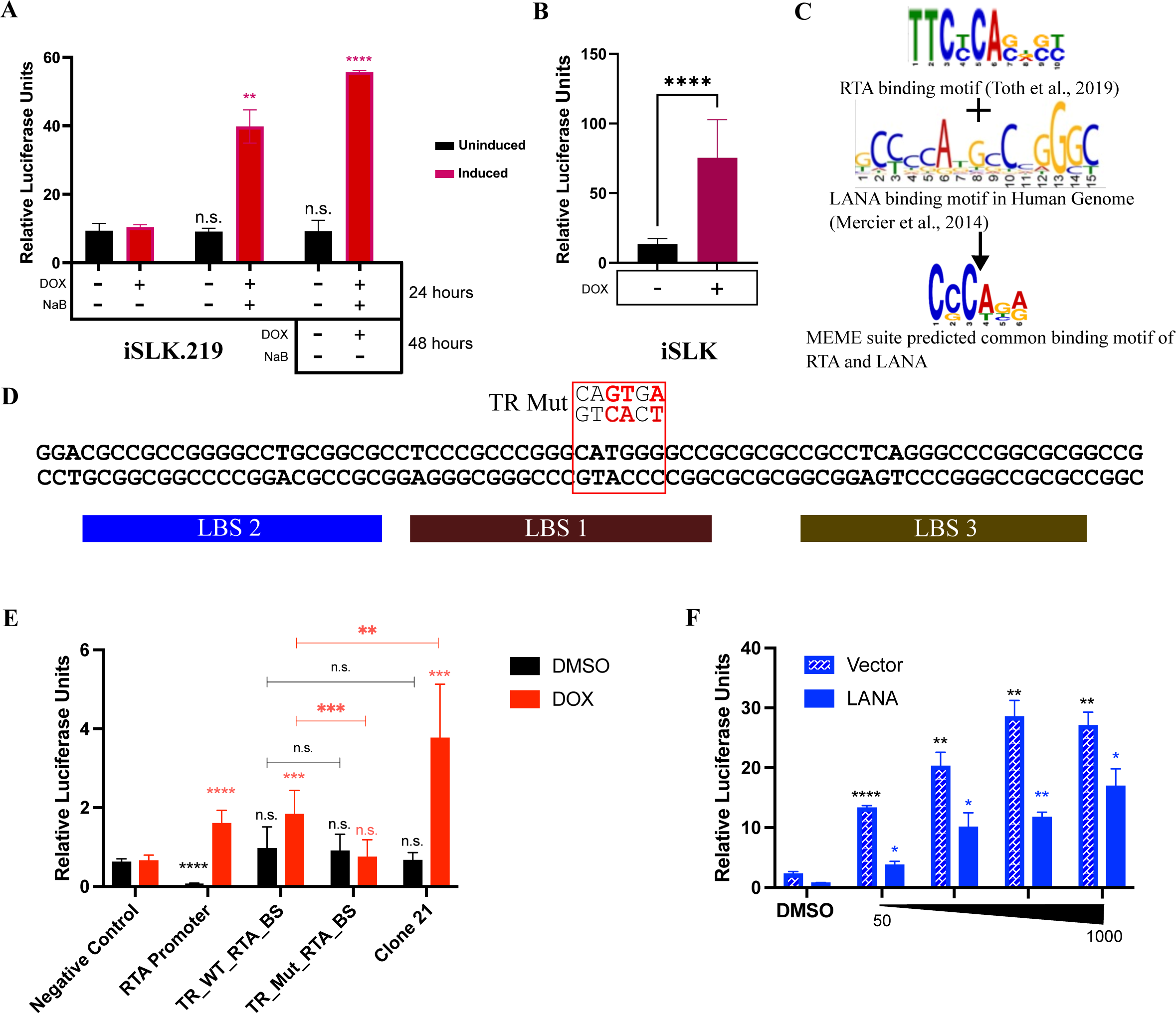
**KSHV Terminal Repeats are RTA responsive**. (A) iSLK.219 cells were transfected with clone 21 and induced with 1μg/ml Doxycycline (Treatment A), 1μg/ml Doxycycline with 1.25mM Sodium Butyrate for 24 hours (Treatment B), or 1μg/ml Doxycycline with 1.25mM Sodium Butyrate for 24 hours followed by only 1μg/ml Doxycycline induction for another 24 hours (Treatment C). (B) iSLK cells were transfected with clone 21 and induced with 1μg/ml Doxycycline (Treatment A), same as in A. (C) Logo binding motifs as revealed by Motif search analysis in MEME suite using RTA responsive viral genomic enhancers, and LANA binding consensus sequence on human chromosome. (D) Sequence of LBS within TR, where LBS1-3 are marked as boxes below the sequence. The MEME-predicted common RTA-LANA binding motif is marked by the red rectangle. Mutated nucleotides used in reporter assay are marked in red above the TR sequence. (E) iSLK cells were transfected with the TR sequence or the mutated as presented in D cloned downstream to the luciferase. Clone 21 (TRx7) served as a positive control. Cells were treated with vehicle (DMSO) or 1μg/ml Doxycycline for 24 h before luciferase assay. (F) iSLK cells were transfected with clone 21 together with a constant amount of LANA expression vector or empty vector and then treated with increasing doses of Dox (50ng, 250ng, 500ng, and 1000ng/ml). Data represent Mean ± S.D. of n=3 biological replicates. *P<0.05; **P<0.01; ***P<0.001; ****P<0.0001.

We hypothesized that LANA represses the TR enhancer, and RTA can activate the TR by over- competing with LANA to gain access to the TR. To test this, we followed TR activity while gradually increasing RTA levels. A luciferase reporter assay with the TR enhancer in iSLK cells, treated with increasing doses of Dox (50ng, 250ng, 500ng, and 1000ng/ml) revealed that RTA can activate the TR regardless of LANA presence (**Fig. 4F**). However, higher levels of Dox were required to achieve the same level of activity in cells expressing LANA. This indicates that RTA can effectively compete with LANA to induce TR enhancer activity.

### Dissecting functional regions within KSHV enhancers

To gain a better understanding of the functional elements of KSHV genomic enhancers, we decided to narrow them down. Primers were designed to amplify the viral enhancers in a manner that enabled us to divide them into different portions: Left (L), Middle (M), and Right (R) (**Fig. 5A** & **Table S2**). Then these truncated enhancers were transfected into iSLK cells.

**Figure 5:**
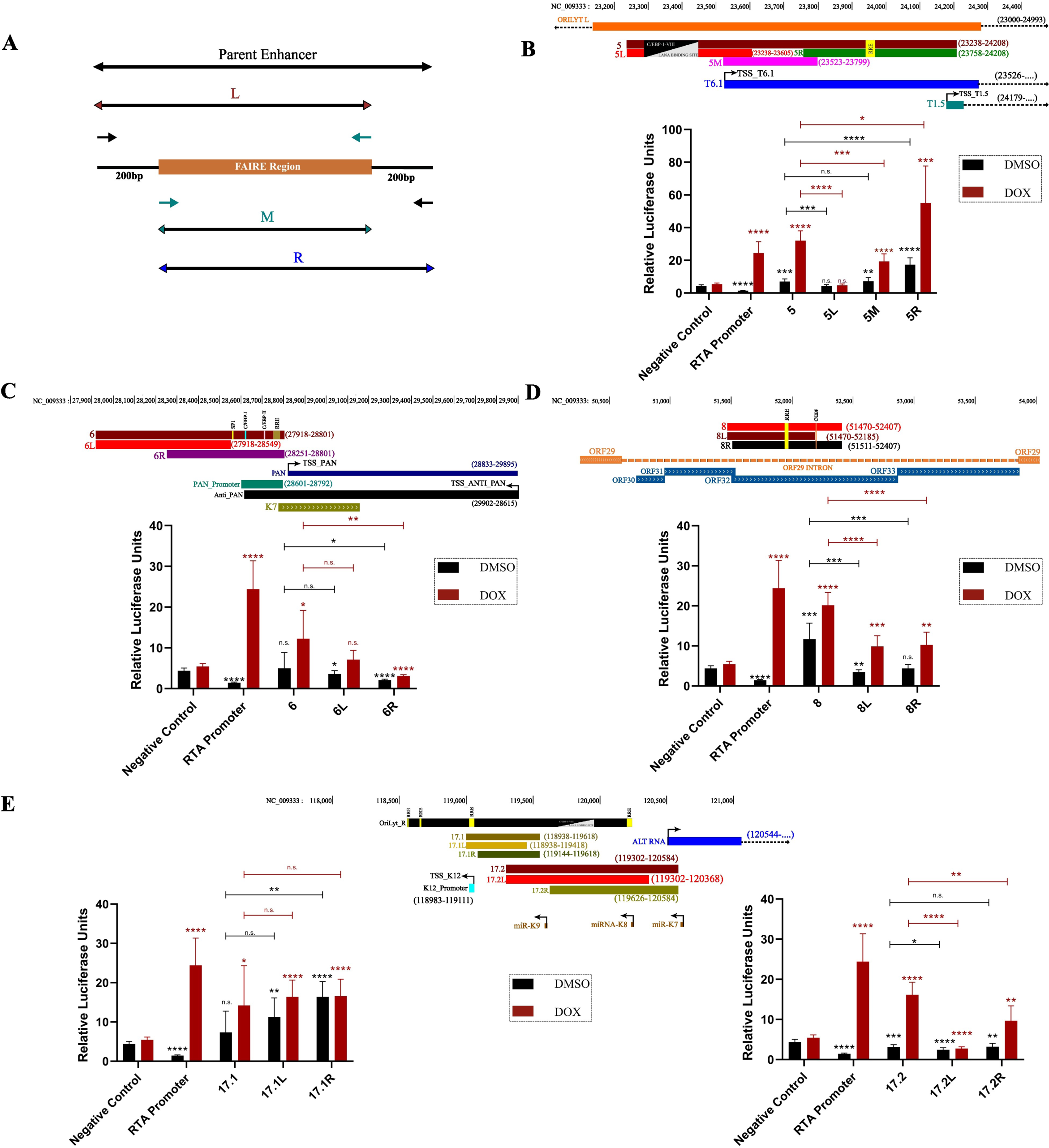
**Dissecting functional regions with KSHV enhancers**. (A) schematic presentation of the strategy to clone the deletions of the identified enhancers. The iSLK cells were transfected with the deletion reporter plasmids for OriLytL (B), PANp (C), ORF29-intron (D), OriLytR-K12 and OriLytR-ALTp (E). Cells were treated with 1μg/ml Doxycycline (red bars) or DMSO (black bars). Data represent Mean ± S.D. of n=3 biological replicates. *P<0.05; **P<0.01; ***P<0.001; ****P<0.0001.

As expected, narrowing down the enhancers affected their activity. The left portion of region 5, denoted as 5L (23238-23605nt), displayed no enhancer activity, whereas the 5M (23523- 23799nt) and 5R (23758–24208) segments maintained their enhancer activities (**Fig. 5B**). Interestingly, both 5M and 5R are associated with two KSHV non-coding RNAs (ncRNAs T6.1 and T1.5). Additionally, 5R contains the RTA Recognition Element (RRE), which likely contributes to its strong RTA-mediated enhancer activity. Surprisingly, region 6 narrow-down resulted in no enhancer activity (**Fig. 5C**), indicating that an intact region 6 is needed for the enhancer activity. Narrowing down region 8 destroyed its constitutive enhancer activity and significantly decreased the RTA inducible activity (**Fig. 5D**). The sequences between 51470- 51511nt seem to be critical for the constitutive activity of ORF29 Intron enhancer. Interestingly, this region harbors a YY1 binding site. The YY1 plays an important role in enhancer activity by promoting enhancer-promoter loop formation (71). Nevertheless, both regions flanking the FAIRE-seq of region 8 are needed for its enhancer activity. Narrowing down region 17.1 did not show any significant reduction in its activity (**Fig. 5E**). However, it became evident that its constitutive nature is attributed to its right portion (17.1R, 119144- 119618). It is noteworthy that region 17.1 consists of K12p and portions of OriLyt-R, which fall within the latency cluster. In the case of region 17.2, narrowing down revealed that only its right portion (17.2R, 119626-120584) associated with ALTp acts as an enhancer. Altogether, the narrow-down experiments revealed the functional regulatory portions of the KSHV enhancer elements.

### KSHV genome contains a Silencer element within its latency cluster

Silencers are distal cis-regulatory elements that repress their target gene promoters through physical interaction in an orientation- and position-independent manner (39, 42, 72). In our FAIRE-STARR reporter screen for KSHV, we identified a silencer region within the viral genome. The 5’UTR of LANA-K14p (clone 19) repressed reporter activity in iSLK (∼5-fold), iSLK.219 (∼3-fold) and HCT116 cells (∼10-fold) (**Fig. 6A, S3**). These results indicate the presence of a viral genomic silencer element in the KSHV latency gene cluster. Interestingly, the repression by this silencer seems to vary between different cell lines, similar to cellular silencers (73–75).

**Figure 6:**
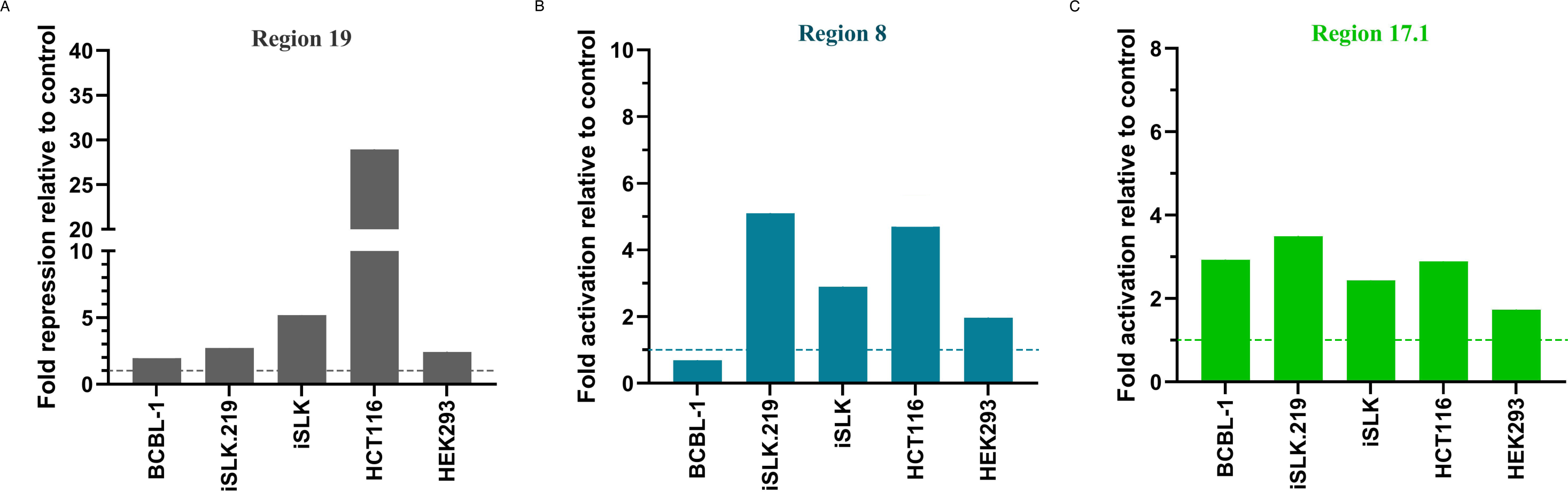
**Tissue-specific enhancer activities**. The silencer in region 19 (A) and the constitutive enhancers in region 8 (B) and 17.1 (C) were transfected into infected B-cell (BCBL1), infected epithelial (iSLK.219) and uninfected epithelial (iSLK, HCT116, HEH293) cells. The results are presented as fold of repression (A) or activation (B, C) by dividing the relative luciferase units (RLU) by the RLU of the negative control.

Furthermore, to determine which portion of this region contributes to the silencing activity, we divided region 19 into 3 portions: 19L (127254-127828nt), 19M (127446-128276nt), and 19R (127900-128474nt). Although all the regions repressed the reporter signal, it was found that 19R contributed the most significant repression (**Fig. S4**). Notably, the reporter activity of 19R was not rescued with RTA induction. In summary, these findings suggest the presence of a silencer element in the LANA-K14 promoter region within the latency cluster of the KSHV genome.

We also determined the tissue specificity of the other two constitutive enhancers, regions 8 and 17.1. While region 17.1 exhibited constitutive enhancer activity in all the tested cell lines, region 8 lost its enhancer activity in BCBL1 B-cells (**Fig. 6B & C**). Therefore, ORF29 Intron (region 8) contains a tissue-specific constitutive enhancer.

### CRISPRa targeted to KSHV enhancers can induce lytic reactivation

The enhancer reporter assays identified several enhancers within the KSHV genome. To test their functional relevance in the context of whole viral transcription regulation, we employed CRISPR activation (CRISPRa) to stimulate their potential transcriptional regulatory activity. SLK.219 cells were transduced with lentiviruses expressing dCas9 that tethers strong activation domain (dCas9-10xSunTag and ScFv-2ERT2-VPH) (51), and specific sgRNAs targeting individual enhancers (**Fig. 7A**). Subsequently, the cells were treated with 4-hydroxytamoxifen to induce CRISPRa activity, and RNA was subjected to sequencing (RNA-seq). This analysis revealed both common as well as gene-specific regulations by KSHV enhancers.

**Figure 7:**
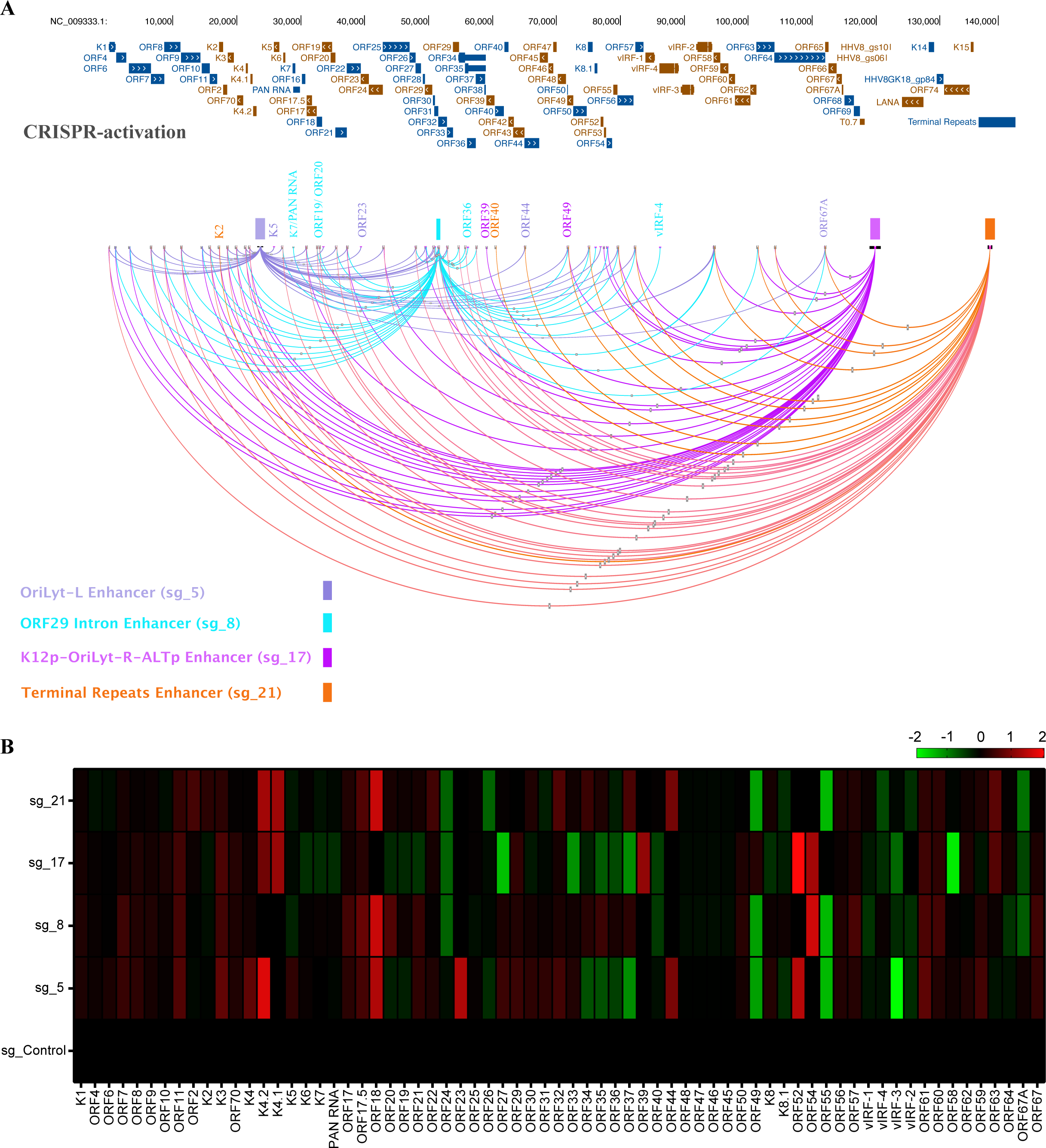
CRISPR activation and RNA-seq reveal genes regulated by viral enhancers. SLK.219 cells were transduced with lentiviruses expressing dCas9 that tethers a strong activation domain (dCas9- 10xSunTag and ScFv-2ERT2-VPH) and specific sgRNAs targeting individual enhancers. Subsequently, the cells were treated with 4-hydroxytamoxifen to induce CRISPRa activity, and RNA was subjected to sequencing (RNA-seq). (A) The affected genes are indicated by a line between the enhancer and the gene. Genes that were strongly regulated by a specific enhancer are indicated with the specific enhancer color. (B) Heatmap presentation of RNA-seq. Gene expression was color-coded relative to sgRNA control. The schema of reference KSHV genome (NC_009333) was generated with UCSC genome browser (54). For each sgRNA, three biological replicates were sequenced.

The CRISPRa approach for the following enhancers, OriLyt-L (region 5), ORF29 Intron (region 8), K12p/OriLyt-R/ALTp (region 17) and TRs (region 21) led to up-regulation of many lytic genes (**Fig. 7**). To validate these findings, we performed the RT-qPCR and confirmed the up-regulation of RTA (ORF 50), k8.1 and K12 (**Fig. S5**). RTA expression levels were upregulated upon performing the CRISPRa of all the enhancers except OriLyt-L (region 5) (**Fig. 7B** and **Fig. S5**). Despite being almost identical, it was interesting to find out that OriLyt-R was able to upregulate the RTA expression levels upon performing CRISPRa whereas OriLyt-L CRISPRa could not. The CRISPRa for OriLyt-L enhancer (region 5) specifically induced the expression of K5, ORF23, ORF44, and ORF67A. The ORF29 Intron enhancer exclusively induced K7, PAN RNA, ORF20, ORF19, ORF36 and vIRF-4 genes. The K12p/OriLyt-R/ALTp enhancer (region 17) specifically induced ORF39 and ORF49; while inducing the Terminal Repeats enhancer (region 21) with CRISPRa led to exclusive upregulation of K2 and ORF40 genes (Fig. 7C).

Although the viral enhancers seemed to induce a significant number of viral genes, performing CRISPRa of any of the viral enhancers led to a specific pattern of gene expression (**Fig. 7B**). It seems that genes and gene clusters were found to be regulated by multiple enhancers. Examples include genes like ORF11, ORF70, ORF50, ORF61, ORF67, etc. Additionally, there were genes regulated by two or three enhancers in combination. For instance, ORF20, ORF19, ORF34-37 appeared to be upregulated by ORF29 Intron and Terminal Repeats, while genes like ORF27, ORF29, ORF30 were induced due to CRISPRa of OriLyt-L, ORF29 Intron and Terminal Repeats (**Fig. 7A & B**). While individual KSHV enhancer elements can regulate specific genes, the results of the CRISPRa experiment indicate that, in most cases, viral enhancers work collectively and complementarily. This suggests that the KSHV transcriptional repertoire is controlled by a collective ‘viral super-enhancer’ hub coming in close contact with each other and with the gene promoters regulated by these viral enhancer elements. This finding also implies that a single enhancer is not solely responsible for controlling the entire lytic transcriptional reactivation. Altogether, our data suggest that viral genomic enhancers possess exclusive, complementary, or collective distal cis-regulatory functions on viral genes or gene clusters both in the latency and lytic phases.

### CRISPRi targeted to KSHV enhancers disrupt the viral transcriptional repertoire

To address the function of KSHV enhancers, we also employed CRISPR interference (CRISPRi) by directing dCas9 fused to a strong transcription repression domain (dCas9- KRAB-MeCP2) (50). SLK.219 cells were transduced with lentiviruses expressing sgRNA and dox-inducible dCas9-KRAB-MeCP2. The cells were then treated with or without TPA in the presence of Dox, and RNA was isolated and subjected to RNA-seq. The CRISPRi of OriLyt-L enhancer (region 5) exclusively downregulated the expression of ORF57, ORF61, ORF62, ORF63, and ORF64 during latency and K4, K5, K6, ORF21 post lytic reactivation (**Fig. 8A- B, S6B-C**). The CRISPRi for PAN promoter (region 6) exclusively downregulated the expression levels of ORF11 and ORF58. Although CRISPRi for ORF29 Intron (region 8) revealed no exclusive gene downregulation in latency, interestingly, marked downregulation of K8 gene was observed during lytic induction. The CRISPRi for K12p/OriLyt-R (region 17) revealed exclusive downregulation of ORF6 and ORF26 genes while for TRs (region 21) resulted in exclusive downregulation of K1 only.

**Figure 8:**
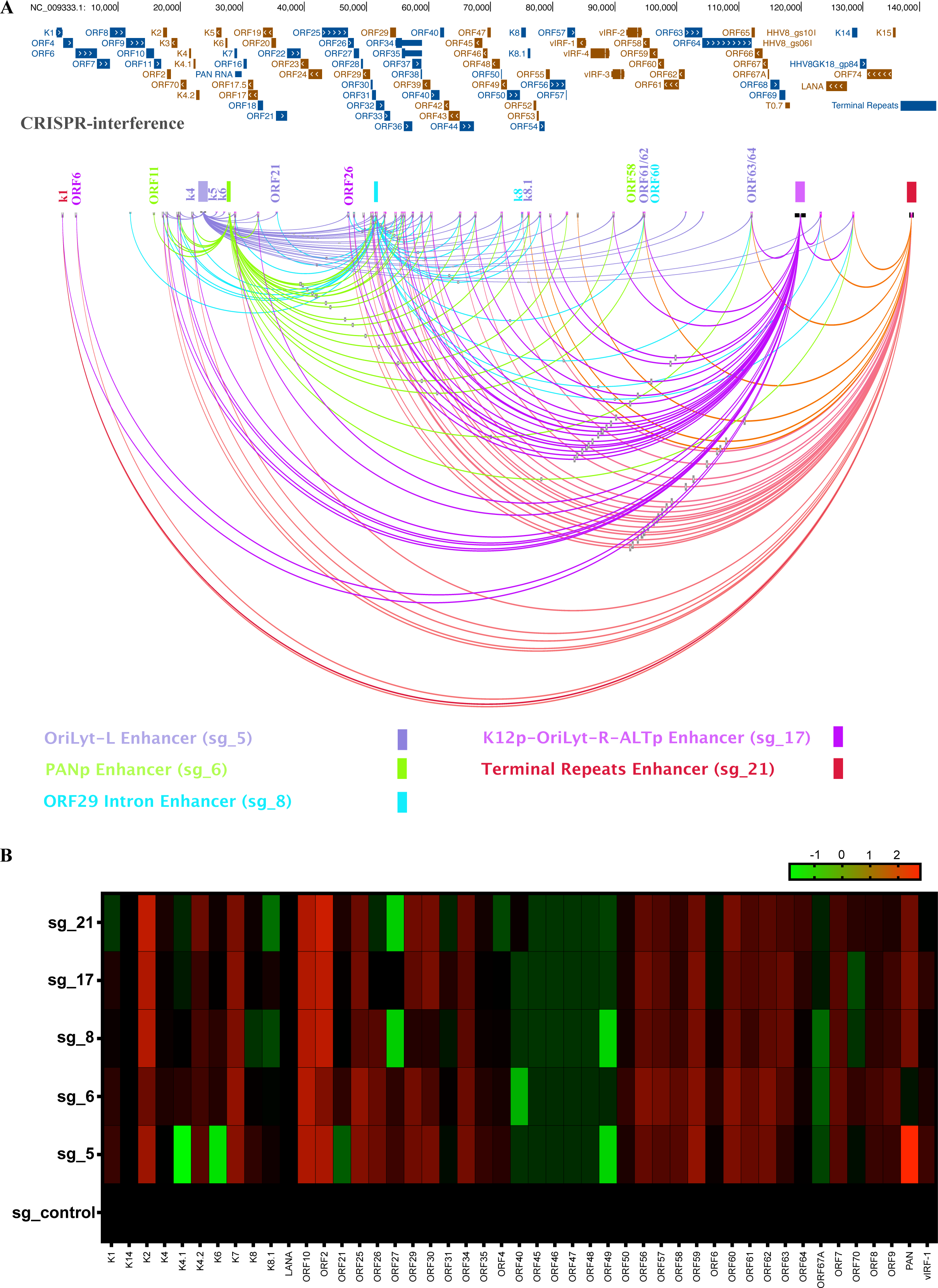
CRISPR interference and RNA-seq reveal genes regulated by viral enhancers. SLK.219 cells were transduced with lentiviruses expressing Dox inducible dCas9 fused to a strong transcription repression domain (dCas9-KRAB-MeCP2) and specific sgRNAs targeting individual enhancers. Subsequently, the cells were treated with TPA in the presence of Dox, and RNA was isolated and subjected to RNA-seq. (A) The affected genes are indicated by a line between the enhancer and the gene. Genes that were strongly regulated by a specific enhancer are indicated with the specific enhancer color. (B) Heatmap presentation of RNA-seq. Gene expression was color-coded relative to sgRNA control. The schema of reference KSHV genome (NC_009333) was generated with UCSC genome browser (54). For each sgRNA, three biological replicates were sequenced.

Surprisingly, we also observed that CRISPRi of any of the viral enhancers downregulated common genes such as K2, K7, ORF18, ORF2, ORF30-38, ORF44-48, and ORF59 (**Fig. 8A- B**). In addition, some genes were controlled by the combination of two or three enhancers. For example, ORF54 was downregulated by CRISPRi for both ORF29 Intron and OriLyt-R; ORF70 was regulated by both PAN promoter and K12p/OriLyt-R, and K8.1 was markedly downregulated by CRISPRi of both ORF29 Intron and TRs during lytic induction. Intriguingly, gene clusters from ORF45-49 seem to be affected collectively by all the enhancers. The CRISPRi experiment supports the notion that viral enhancers work in a collective and complementary manner, in addition to their gene-specific regulation.

## Discussion

The identification of enhancers within the cell genome can be carried out based on their characteristics, such as open/accessible chromatin, specific histone marks, and enhancer- promoter contacts. While, in many cases, specific histone marks such as H3K27Ac, H3K4Me1, and H3K4Me3 can distinguish and predict enhancers and promoters (44, 76), recent research indicates that the function of these two entities may not always be distinct. Promoters can act as enhancers for other genes (77), and enhancers frequently give rise to transcripts (78). This distinction based on histone marks becomes even more challenging in compact viral genomes where every genomic sequence has multiple functions. Enhancer-promoter contacts can be identified through chromosome conformation capture (3C) and its genome-wide versions (79, 80), Hi-C (81) and HiChIP (82). Several studies revealed these interactions along the KSHV genome (58, 83); however, when two promoters are in close proximity, it cannot predict which is solely a promoter and which is a promoter that can also serve as an enhancer. In any case, these previously identified interactions are in agreement with the enhancers and their responsive promoters that we identified in this study (as will be discussed below). Another characteristic feature of enhancers is their open chromatin due to the lack or redistribution of nucleosomes. To detect open/accessible chromatin regions, several methods have been developed, such as DNaseI hypersensitivity-seq (DNase-seq) (84–86), MNase-seq (87, 88), ATAC-seq (89), and FAIRE-seq (47). To identify open chromatin regions in the KSHV genome, Hilton et al. (49) performed FAIRE-seq and identified 24 such regions. Because FAIRE-seq regions are good predictors of enhancers (48), we decided to use these regions in our functional enhancer assay.

Functional enhancer assays rely on the unique ability of enhancers to enhance transcription when they are located downstream of a gene promoter. STARR-seq is a robust method for identifying functional enhancer elements, where putative enhancers are cloned downstream to a weak promoter that transcribes a reporter gene or ORF (55). To systematically search for enhancers and silencers in the KSHV genome, we adopted a combined approach using FAIRE-seq regions in STARR-seq assay. Because of the small number of FAIRE-seq regions in the KSHV genome, we skipped the sequencing step and cloned all regions in the luciferase validation vector. The 21 open chromatin regions identified in the KSHV genome by FAIRE-seq were cloned downstream to the luciferase gene in a STARR-seq luciferase reporter plasmid to make the KSHV-FAIRE-STARR library. The reporter assay for the KSHV-FAIRE-STARR library revealed enhancer activity associated with OriLyt-L, PAN promoter, ORF29 Intron, K12 promoter/OriLyt-R/ALT promoter, and the Terminal Repeats.

Our finding that KSHV harbors several cis-regulatory elements, such as enhancers and silencers (**Fig. 9**), aligns with findings in other viruses. The enhancement of a promoter by a DNA sequence in cis led to the discovery of the first enhancer in the genome of Simian virus 40 (SV40) (90). Later, an enhancer in the Hepatitis B Virus (HBV) genome was identified (91). Interestingly, even the small HBV genome of 3,200 bp, contains more than one enhancer (92). Enhancers are also prevalent in retroviruses, including human immunodeficiency virus (HIV), and human T cell lymphotropic virus (HTLV) (93–95). Viral enhancers have been identified in herpesviruses as well. For instance, the herpes simplex virus type 1 (HSV-1) immediate early (IE) mRNA 3 (96) and the latency-associated transcript (LAT) (97) serve as lytic and latent enhancers, respectively. A strong enhancer is located upstream of the major immediate-early gene of Human Cytomegalovirus (HCMV) (98). Additional enhancers in HCMV are located in the short unique region (99) and OriLyt (100). Even the closest relative of KSHV, the Epstein Barr Virus (EBV), contains several enhancers, including the DR enhancer, which is both constitutive and lytically induced (101). The family of repeats within the oriP is activated by EBNA1, serving as a transcriptional enhancer (102), and the BamHI-C fragment of EBV, responsible for the expression of EBNA-1 in some cell lines, acts as a B-cell-specific enhancer (103). Moreover, there are two TPA-responsive enhancers, MSTRE-I and MSTRE-II, in the upstream sequence of the MS gene of EBV (104).

**Figure 9:**
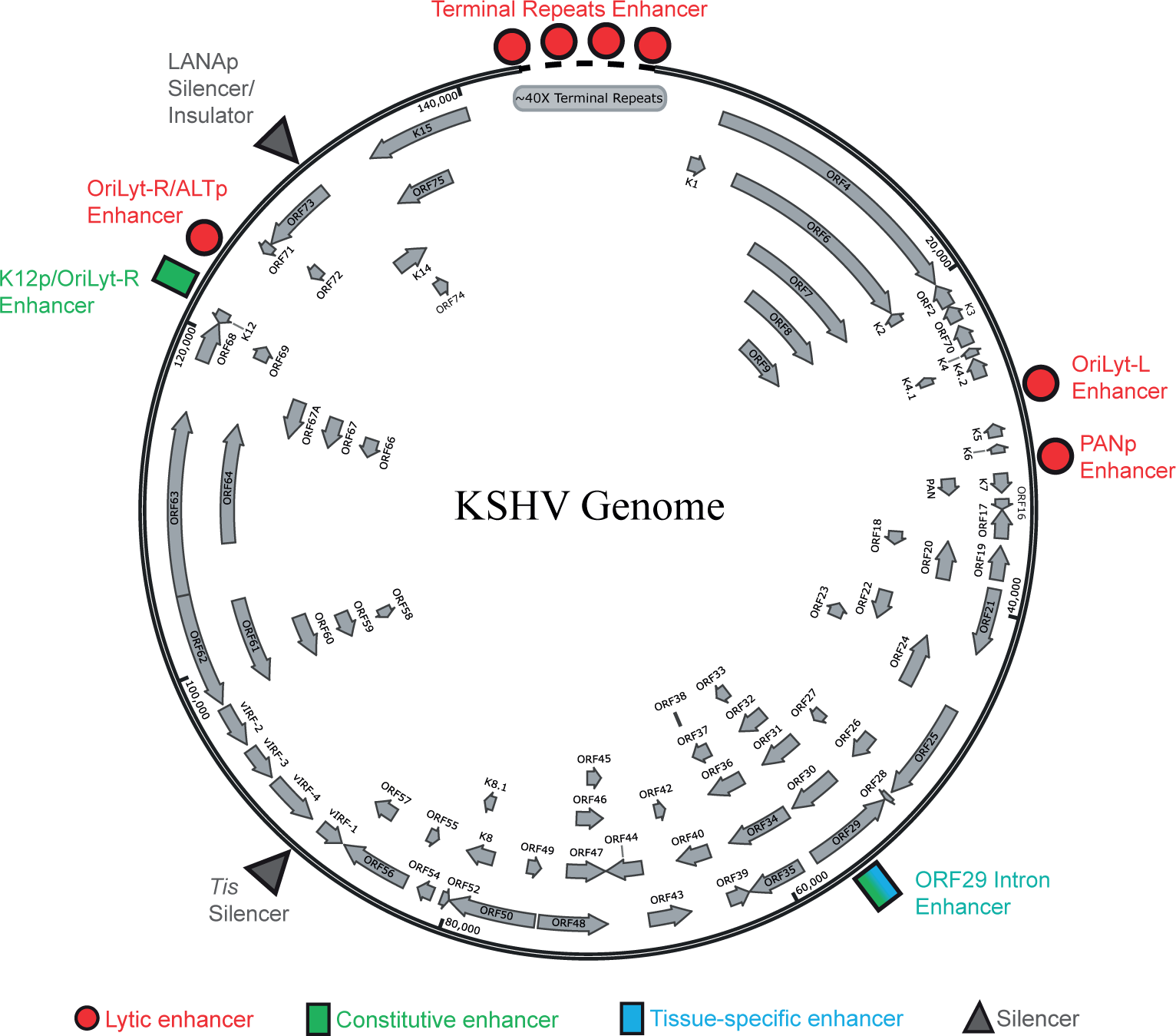
Schematic map of the KSHV enhancers. Schematic map of the KSHV genome that includes the constitutive and lytic enhancers within the genome. Induced/lytic enhancers are marked as red circles, constitutive enhancer is marked by a green rectangle, and tissue-specific enhancer is marked by a blue rectangle. In addition, the latency locus LANAp silencer that might function as an insulator, and the previously (ref) identified Tis silencer are marked by grey triangles. The KSHV genome schema with the ORFs was generated with SnapGene software (www.snapgene.com, v7.0.2).

Since enhancers play a crucial role in tissue-specific gene regulation, we performed luciferase assays in various cell lines, both uninfected and infected with KSHV. The induction of the lytic phase resulted in enhancer activity for OriLyt-L, PANp, OriLyt-R/ALTp and TRs. While K12p functioned as a constitutive enhancer, its activity increased upon lytic induction. Notably, ORF50 Intron (Region 13) and vIRF1 (region 14) also showed enhancer activity in iSLK cells but failed to do so in HCT116 cells. The HCT116 cells, lacking an IFN-mediated response upon plasmid transfection, are utilized in STARR-seq to exclude false positives. This observation suggests that the enhancer activity of these two regions was actually a false positive, and thus, we decided not to proceed with these regions for any further experiments. The IFN-mediated response of vIRF1 (region 14) might be important during the early stages of viral infection. This region encodes proteins that aid the virus to combat cellular interferon response upon viral entry. However, following the establishment of latency and expression of other viral inhibitors of interferon, this region is no longer needed and becomes repressed, as can be seen in latently KSHV-infected cells (iSLK.219 and BCBL-1).

Analyzing enhancer activity in different cell lines revealed that ORF29 intron is a tissue-specific enhancer active in epithelial cells but not in B-cells. Considering the important role of KSHV-encoded RTA in the regulation of lytic gene expression and its binding, as determined by ChIP-seq (57, 58), to all the lytic enhancers we identified, we explored its role in enhancer regulation. Using both uninfected cells with dox inducible RTA expression and transient transfection, we found that the KSHV lytic enhancers, OriLyt-L, PANp, OriLyt-R/ALTp and TRs, are all RTA responsive. Interestingly, the EBV functional homolog of RTA, ZTA (also known as BZLF1, Z, and EB1), also activates EBV enhancers (105).

KSHV possesses two lytic replication origins (OriLyts) in its genome (23, 106). Our study indicates that OriLyt-L functions as a potent lytic enhancer. Within the OriLyt-R region, there are two enhancers: one constitutive and one lytically induced. This entire region is active during latency and is further induced by RTA during the lytic cycle (107–109). The K12 promoter is flanked by OriLyt-R (108). This region is also known to transcribe several miRNA, expressed during latency (110–114). The presence of these enhancers in the latency cluster and the constitutive nature of K12p/OriLyt-R enhancer suggests their potential regulatory role on latent gene expression. Interestingly, the OriLyt-R/ALTp enhancer region is only active during the lytic phase in contrast to its flanking counterpart K12p/OriLyt-R. The OriLyt-R has been suggested to possess long-range transcriptional regulation properties based on loop formation with RTA promoter and the observation that a mutation in the RTA binding site within OriLyt-R abrogated RTA expression during exogenous RTA lytic induction (58). HCMV and EBV OriLyts have been previously reported to exhibit enhancer activities (100, 106). OriLyts in herpesviruses are somewhat conserved, and KSHV OriLyts also contain the C/EBP CCCAT domains, supporting our finding of OriLyt-L as an RTA-inducible enhancer (115). Additionally, similar to KSHV, HCMV OriLyt gives rise to two ncRNAs (100). Thus, it can be envisioned as an enhancer, giving rise to ‘enhancer-derived lncRNA’ (116–118). In summary, our observation that KSHV OriLyts function as enhancers aligns with findings in other herpesviruses.

Another strong lytic enhancer identified in our study is the PAN promoter. The PAN promoter possesses a Sp1 binding site, along with some human genomic enhancer-like sites in its sequence (119). The utilization of the enhancer element within the PAN promoter region seems to be another way the virus exploits host cell machinery for its benefit. Indeed, the PAN promoter serves as an enhancer in all the cell lines used in this study. A previous study detected strong interaction/looping between PAN promoter and RTA promoter (58). To further explore this interaction, we cloned PANp downstream to RTAp, and vice-versa, and performed functional enhancer assays. This set of experiments clearly shows that PANp can function both as a promoter and an enhancer that can enhance transcription from RTAp. In contrast, RTAp can only serve as a promoter. Therefore, our study determined the functional relations between the previously identified interactions, providing evidence that PANp is a lytically induced enhancer.

Enhancers might reside within introns (32, 56). The ORF29 Intron resides in the antisense strand of ORF32 (108). To date, it was unclear the exact functional relevance of this intron sequence despite the fact that this particular stretch of the viral DNA harbors CTCF and RTA binding sites, all when the ORFs associated with this intron are late lytic in nature (13, 49, 77). In our in-silico motif enrichment analysis of the ORF29 Intron, a YY1 binding site (CCAT) was revealed. YY1 is well known for facilitating specific enhancer-promoter chromatin looping (71). While reporter assays in BCBL-1 did not show any enhancer activity for this region (region 8), enhancer activity was observed in epithelial-origin cell lines like iSLK and HCT, even in the absence of RTA expression. This observation indicates its role as a constitutive cell-type/tissue-specific enhancer.

The KSHV genome contains around 30 copies on average (∼16-75 copies) of terminal repeat (TR) sequences (10, 120). Each repeat is a ∼801bp of GC-rich region that is an open chromatin region and harbors CTCF binding sites (13, 49). GC-rich sequences often serve as enhancers and attract multiple transcription factors (121, 122). Garber et al., 2002 (10) reported that TR cloned downstream to SV40 promoter can enhance transcription in HEK293 cells and suggested that it possesses enhancer activity. Moreover, they have found that this enhancer activity is strongly suppressed by LANA. A recent study (123) performed both proteomic and Cleavage Under Targets & Release Using Nuclease (CUT & RUN) for RTA and LANA, revealed shared protein complexes between LANA and RTA at the TR. Similar to our study, they suggest that the TR is an RTA inducible enhancer. Our observation that TR enhancer activity is reduced in infected cells compared to uninfected cells expressing only RTA, is in agreement with LANA’s ability to repress the TR. In infected cells, Dox was not sufficient, and treatment with TPA and SB in addition to Dox to achieve lytic induction and higher concentrations of RTA were needed. By using uninfected cells expressing only LANA and increasing concentrations of RTA, we were able to show that RTA was able to outcompete LANA on TR activity without the need for any other viral factor. We identified an RTA binding motif within LBS1, and mutation at this site abrogated the ability of RTA to enhance TR activity. The fact that LANA represses the TR and, therefore, higher concentrations of RTA are needed to activate this enhancer suggests that LANA safeguards latency, ensuring the lytic cycle is turned on only when a sufficient amount of RTA is expressed. This role is in addition and in agreement with its role in repressing the RTA promoter (27–29). The repetitive nature of the TR makes it an even stronger enhancer. We have found that one repeat already can serve as an enhancer, and seven copies of TR were the strongest enhancer in our functional enhancer assay. However, the KSHV genome harbors about 30 copies of TRs, making it a super-enhancer. Cellular super-enhancers are characterized by large size, high abundance of transcription factors, high H3K27Ac and BRD2/BRD4, and sensitivity to perturbations (124). The TR is large, spanning around 24 kbp, highly abundant in H3K27Ac (61, 125) and its reader BRD2 and BRD4 (126). It is tempting to speculate that, similar to super-enhancers, it is sensitive to perturbations.

We also came across a silencer element in the KSHV genome. Silencers are cis-regulatory elements that repress their target gene promoters by recruiting repressor proteins (39, 127). These elements are also responsible for tissue/cell-specific gene expression (38). Silencers are often found within various sites like introns, gene promoters, UTR regions, etc (38, 127, 128). We found that a region upstream to the LANA promoter (region 19) repressed the reporter signal by ∼3-10 fold in different cell lines, where the lowest repression was observed in BCBL-1. This indicates the silencer activity to be tissue-specific. The presence of activating H3K4Me3 marks in LANA-K14p (49) does not mean that it cannot act as a genetic silencer, as H3K4Me3 marks often colocalize with the repressive H3K27Me3 marks in 75% of instances (128). To find out the core silencer element, we narrowed down the silencer like the viral enhancers. This revealed that the sequence spanning 127900-128474 nt contributed most to the silencer activity of the region. This region is located between the promoter for the latency genes LANA/v-FLIP/v-Cyclin and the promoter for the lytic genes ORFK14 and ORF74. This region has been shown to be bound by CTCF, RAD21 and SMC1/SMC3 (129, 130). Cohesins consist of four major subunits, SMC1, SMC3, RAD21/Scc1, and Scc3, which have been shown to form a ring-like structure involved in sister-chromatid cohesion, organization of topologically associated domains, and enhancer-promoter interactions. CTCF is associated with cohesin and directs the sites of cohesin on the chromatin (131). The binding of CTCF at the human imprinted region *h19/igf2* indicated its role as an enhancer blocker or insulator (132). Insulators are CTCF-binding sites that can prevent enhancers’ effect on promoters. Deletion of the CTCF sites within this LANA promoter region, as well as knock-down of RAD21, led to specific up- regulation of the lytic genes ORFK14 and ORF74 (129), indicating a role of an insulator to this region. Therefore, this region may serve a dual role as a silencer and insulator. Previously, a silencer was identified upstream of the ORF K9 (vIRF1) gene and termed *Tis* (133). So, it seems that the KSHV genome harbors at least two silencers. The related herpesvirus HCMV harbors a silencer in a region containing 7R1 that is able to down-regulate transcription from either the US3 promoter or a heterologous promoter in a position- and orientation-independent manner (134). Silencers were also identified in the genome of EBV (104).

To reveal the functional aspects of the KSHV enhancers, they were targeted by CRISPR activation (CRISPRa) and interference/repression (CRISPRi), and total RNA sequencing was performed. This revealed both gene-specific regulation, as well as global and common regulation by several enhancers. This observation indicates that KSHV enhancers not only have specific but also collective regulatory activity on viral gene promoters. Also, this suggests that KSHV has a ‘viral super-enhancer hub’ to control its transcriptional network. The genes that were repressed due to viral enhancer CRISPRi, were in line with the findings of CRISPRa, where the same genes were upregulated (Fig. 7A & 8A). This not only validated the CRISPRi result, but also proved the relevance of the viral enhancers activity during their transcriptional repression and activation, respectively. The CRISPRa and CRISPRi data were also in agreement with the KSHV capture HiC data published previously (58). Compellingly, we found that transcriptional activation of most of the viral enhancers upregulated RTA (ORF50), a critical factor responsible for viral lytic reactivation. This indicates the cooperation among the viral enhancers to induce the lytic reactivation. Unexpectedly, we found that CRISPRi to the enhancers also led to RTA upregulation. Not only that but many other late lytic genes were also found to be upregulated under this condition. So why did CRISPRi to KSHV enhancers led to the upregulation of RTA? Since the KSHV genome is small and compact relative to the human genome, any perturbation of the viral genome with CRISPRi would lead to the disruption of the normal 3D conformation of the viral genome. This perturbation of the viral enhancers would lead to wide-scale conformation and transcriptional disruption. This disruption would help the transcriptionally inactive RTA promoter to come into the vicinity of the rest of the viral enhancers which are already enriched with active histone marks (like H3K27Ac). One might ask how it is possible for a lytic enhancer to induce the RTA promoter without the RTA expression itself. Our observation that PANp enhancer when located downstream to RTAp can enhance RTAp activity even without RTA expression supports this model. Several studies have suggested that the KSHV genome is folded in two different ways one as latent and second as lytic (83, 135). Therefore, targeting dCas9 may disrupt this latency-specific folding, leading to uncontrolled lytic expression. The same effect on lytic gene expression was observed when dCas9 was targeted to EBV enhancer (136).

In summary, the systematic functional analysis presented here identified several enhancers in the KSHV genome and their relevance to the KSHV transcriptional program. Enhancers are distal cis-regulatory elements that hold a key role in transcription regulation. It will be interesting to test in the future their contribution to viral gene expression under different physiological and cell differentiation conditions and their roles in directing latent and lytic phases. This new enhancer and silencer map gives us a better understanding of the complex regulation within the KSHV genome and may be exploited in the future for viral therapy.

## Data availability

Large data files will be deposited to NCBI GEO upon acceptance of the manuscript.

## Funding

This work was supported by grants from the Israel Science Foundation (https://www.isf.org.il) to M.S. (1365/21), and the Israel Cancer Research Fund (https://www.icrfonline.org/) to M.S. (23-101-PG). We are grateful for the support of the Elias, Genevieve and Georgianna Atol Charitable Trust to the Daniella Lee Casper Laboratory in Viral Oncology. The funders had no role in study design, data collection and analysis, decision to publish, or preparation of the manuscript.

## Acknowledgments

We would like to thank Alexander Stark, Rolf Renne, Andrea Califano, Yu Wang , Ron Vale, Didier Trono, and S. Diane Hayward for kindly providing plasmids, and Don Ganem, Rolf Renne, Jeffrey Vieira, and Richard F. Ambinder, for cell lines. We would like to thank Shira Perez and Marina Kurtz for library preparation and RNA sequencing and Ronit Sarid for critically reading this manuscript.

